# Cytologic Scoring of Equine Exercise-Induced Pulmonary Hemorrhage (EIPH): Performance of Human Experts and a Deep Learning-Based Algorithm

**DOI:** 10.1101/2022.02.28.482322

**Authors:** Christof A. Bertram, Christian Marzahl, Alexander Bartel, Jason Stayt, Federico Bonsembiante, Janet Beeler-Marfisi, Ann K. Barton, Ginevra Brocca, Maria E. Gelain, Agnes Gläsel, Kelly du Preez, Kristina Weiler, Christiane Weissenbacher-Lang, Katharina Breininger, Marc Aubreville, Andreas Maier, Robert Klopfleisch, Jenny Hill

**Author notes:** CAB, CM and AB have contributed equally.

## Abstract

Exercise-induced pulmonary hemorrhage (EIPH) is a relevant respiratory disease in sport horses which can be diagnosed by examination of bronchoalveolar lavage fluid (BALF) cells using the total hemosiderin score (THS). The aim of this study was to evaluate the diagnostic accuracy and reproducibility of trained annotators and to validate a deep learning-based algorithm for the THS. Digitized, iron-stained cytological specimens were prepared from 52 equine BALF samples. Ten annotators produced a THS for each slide according to published methods. The reference methods for comparing annotator’s and algorithmic performance included a ground truth dataset, the mean annotators’ THSs, and chemical iron measurements. Results of the study showed that annotators had marked inter-observer variability of the THS, which was mostly due to a systematic error between annotators in grading the intracytoplasmatic hemosiderin content of individual macrophages. Regarding overall measurement error between the annotators, 87.7% of the variance of the could be reduced by using standardized grades based on the ground truth. The algorithm was highly consistent with the ground truth in assigning hemosiderin grades. Compared to the ground truth THS, annotators had an accuracy of diagnosing EIPH (THS of < or ≥ 75) of 75.7% whereas the algorithm had an accuracy of 92.3% with no marked difference in correlation to chemical iron measurements. The results show that deep learning-based algorithms are useful for improving reproducibility and routine applicability of the THS. For THS by experts, a diagnostic uncertainty interval of 40 to 110 is proposed. THSs within this interval have insufficient reproducibility regarding the EIPH diagnosis.

## Introduction

Exercise-induced pulmonary hemorrhage (EIPH) in horses is a disease characterized by (repeated) hemorrhage from the lungs during high-intensity athletic activity.^15^ This disease is reported with very high prevalence in numerous breeds of sport horses.^33^ Although the underlying pathophysiological mechanisms and predisposing risk factors of EIPH are not fully understood, it has been shown that severe EIPH has a negative impact on athletic performance in horses.^10,15,16^

Following pulmonary bleeding, red blood cells (RBC) are removed by mucociliary clearance through the upper airways or degraded to hemosiderin (iron-protein-complex derived from breakdown of hemoglobin) by alveolar macrophages. The presence and severity of EIPH can be evaluated by tracheobronchoscopic examination,^10,15^ or quantification of RBC components in respiratory tract fluid. Although there are numerous diagnostic methods with different specificity and sensitivity, noted below, a true gold standard method is lacking.^9,11,34^

Tracheobronchoscopic evaluation of blood content in the airways shortly after strenuous exercise has been proposed as the best available method by the American College of Veterinary Internal Medicine.^15^ This method is relatively easy to perform and seems to have a very high specificity.^18^ However, sensitivity was estimated to be only 59% (many false negative diagnoses) when compared to RBC content in respiratory fluid cells.^18^ Therefore, it has been proposed that a lack of tracheobronchoscopic evidence of blood cannot be used to rule out EIPH.^18,28^

Diagnosis of EIPH through examination of respiratory tract fluids has been derived from the RBC content,^18,30^ hemosiderin content in alveolar macrophages,^11,13^ or less commonly hemoglobin concentration.^30^ As compared to tracheobronchoscopy, those tests are generally assigned a higher sensitivity and many authors have recommend these as the best available diagnostic tests.^13,16,18,28,33,34^ Whereas RBC counts can only be used to diagnose a recent EIPH episode within a few hours to days, increased hemosiderin content in alveolar macrophages (i.e. hemosiderophages) may reveal less recent EIPH episodes.^9^ Previous studies have found that increased numbers of hemosiderophages can be detected from day 7 up to 28 days after a single event of pulmonary bleeding or experimental blood inoculation.^24,28,32^

Different cytologic, semi-quantitative scoring systems to evaluate hemosiderin content in alveolar macrophages have been proposed, which either use conventional cytological staining or specific iron stains (e.g. prussian blue) for hemosiderin.^11-13,17,29^ The most complex scoring system by Doucet and Viel (2002) grades the intracytoplasmic hemosiderin content of 300 alveolar macrophages into 5 tiers, based on the amount of blue hemosiderin pigment, using special iron stain. Scoring ranges from 0 (absence of intracytoplasmic hemosiderin) to 4 where macrophages are filled with hemosiderin. Subsequently, the Total Hemosiderin Score (THS) is calculated per 100 cells, and can range from 0 – 400. Compared to post-exercise tracheobronchoscopy, the presence of EIPH was best predicted at a cut-off value of THS ≥ 75, with a sensitivity of 94% and a specificity of 88%.^13^ Although this scoring system is probably the most sensitive and presumably most reproducible diagnostic test currently available, it has been declared unsuitable for routine diagnostic use due to the high expenditure of human labor.^9^ Regardless of its quantitative nature, previous studies have also shown that grading hemosiderin content of individual cells based on the definition by Doucet and Viel (2002) has some inter- and intra-rater inconsistency.^20,21^

In order to overcome these limitations of the THS, a deep learning-based algorithm for automated image analysis has been recently developed by our research group.^20,22^ Automated image analysis is a field of great interest in veterinary medicine and is becoming increasingly feasible with incorporation of Whole Slide Image (WSI) scanners into the workflow of veterinary laboratories, appropriate IT infrastructure and computational power, and advancing artificial intelligence methods, specifically deep learning.^6,23,27,35^ However, a thorough validation of those algorithms is necessary before they can be used for routine diagnostic service or clinical research.^23,27^

The aim of the present study was to determine the inter-observer variability of the THS and to validate the performance of a deep learning-based algorithm compared to ten human experts, a ground truth dataset and chemical measurements. Our hypothesis was that the use of a deep learning-based algorithm allows an efficient THS analysis while having a high diagnostic consistency and accuracy.

## Material and Methods

### Study specimens (cytologic whole-slide images)

For this study, 29 different bronchoalveolar lavage fluid (BALF) samples from 25 horses, including two samples from each of four horses with separate BALF samples from the left and right lungs, were prospectively collected from routine diagnostic samples submitted to VetPath Laboratory Services (Ascot, Australia). Twenty-eight samples were submitted for routine evaluation of EIPH and one case was submitted for routine evaluation of equine asthma. Use of these samples for this study was approved by the State Office of Health and Social Affairs of Berlin, Germany, approval ID: StN 011/20. Two cytological specimens per BALF sample were prepared using cytocentrifugation (CYTOPRO 7620, Wescor Inc, Logan, UT, USA) of a variable volume of BALF (depending on cellular density) at 510 x g for 3 minutes. Unstained specimens were sent to the FU Berlin, Germany, and one of the two specimens was stained with Perl’s Prussian blue and the other using a modified Turnbull’s blue reaction, Quincke reaction, according to standard protocols.^31^ In both cytochemical staining methods, nonheme-iron reacts with the staining solution forming an insoluble blue pigment.^25^ Hemosiderin is largely composed of ferric iron, Fe^+3^; however, there is also some ferrous iron (Fe^2+^) present along the margins of hemosiderin.^26^ While Prussian blue stains iron in the ferric state, Turnbull’s blue detects ferrous iron and is considered less suitable to stain hemosiderophages. However, the Quincke reaction uses a pretreatment with ammonium sulfide that reduces ferric iron to ferrous iron, therefore iron of both oxidation states is stained by the modified Turnbull’s blue reaction.^31^ Nuclear Fast Red solution was used to counterstain nuclei. Although previous studies on equine EIPH mainly used Prussian blue,^11,13^ we used both staining methods in an attempt to increase image variability which in turn might improve the robustness of the developed algorithm. All slides were digitized with a linear scanner (ScanScope CS2; Leica) in one focal plane at 400x magnification and a resolution of 0.25 µm per pixel. Focus points for scanning had to be selected manually for some slides in order to improve WSI quality. One slide stained with Prussian blue was excluded due to insufficient number of cells (< 300) present on the slide. In 12 WSIs stained with the modified Turnbull’s blue method, budding fungal hyphae and conidiophores were detected. The most reasonable explanation was fungal contamination of the staining solution. As fungal conidiophores may be difficult to distinguish from hemosiderophages with special iron staining, we excluded two WSIs with a proportion of >1% among alveolar macrophages. The other 10 cases had a proportion of conidiophores of <<1% and influence on the THS was considered negligible. Of the three excluded WSIs, the corresponding WSI of the same BAL sample with the other staining method was also excluded for consistent statistical analysis. The final study set comprised 26 WSIs for each staining method, i.e. a total of 52 WSIs.

### Annotators’ scoring

THSs of the 52 WSIs were performed by ten annotators (JS, FB, JBM, AKB, GB, MEG, AG, KdP, KW, JH) including 5 board-certified veterinary clinical pathologists, 4 veterinary clinical or anatomic pathologists in training and one equine internal medicine specialist with good experience in equine BALF cytology. Participants were provided with the original publication on the THS system for horses ^13^ and were instructed to follow that method. No further instruction or specification was given regarding the application of the scoring system. To both view WSIs and label each cell included in the THS, the offline SlideRunner annotation software ^3^ (N = 9) or the online annotation platform EXACT ^19^ (N = 1) was used. Each annotators created a database containing centroid coordinate annotations of the enumerated macrophages with individual label classes for the five hemosiderin grades. In order to ensure that annotators labeled at least 300 cells, we developed a plug-in software tool that automatically counted the annotations of all label classes combined and notified the annotator when 300 annotations had been made. Seven annotators measured the time required to perform the THS in each WSI. Time measurement started with labeling the first macrophage and ended after labeling the last macrophage.

### Supervised deep learning-based algorithms

For development of deep learning-based models using supervised learning, a state-of-the-art object detection network, RetinaNet, was used as previously described by Marzahl and colleagues.^20^ The model was trained with reference annotations (ground truth dataset) for the 52 cases (see below). The cases of the ground truth dataset were split into three groups for three-fold cross validation. Three different models were developed that each used a different subset of the data for training of the model (training set), validation of the training process (validation set) and testing the performance of the final model (test set). Thereby, we were able to analyse all 52 WSIs with our algorithms while avoiding testing algorithmic performance on the same images that were used for training or validation. To guarantee that all three subsets of the split dataset contained cases with grade 4 cells, we sorted the cases by their number of grade 4 cells and assigned them in alternating order to these groups. The models were trained with the Adam optimizer and a maximal learning rate schedule of 0.001 until convergence was reached on the respective validation set (early stopping paradigm) as previoulsy described.^20^

### Reference methods

A true gold standard for quantification of hemosiderin in BALF is not available.^9,11,34^ For comparison of the performance of the THS determined by 10 annotators and the deep learning-based algorithm, we used different reference methods: 1) mean annotators’ THS, 2) ground truth THS, 3) laboratory tests (RBC count, hemoglobin and iron concentration). The human and algorithmic performance in assigning individual macrophages into the five hemosiderin grades was compared to the ground truth cell annotations.

### Mean annotators’ THS

The mean THS was calculated for each WSI based on the ≥300 annotations of each annotator (see above). This reflects the consensus of the 10 annotators and thereby the systematic error between each annotator is averaged.

### Ground truth annotations

The ground truth (a theoretical concept of ‘correct’ annotations) dataset used in this study contained annotations for all alveolar macrophages of the 52 WSIs. Annotations were created by one experienced annotator (CB) using a computer-assisted labeling approach in order to reduce human error. For the analysis of the study results, either labels of the hemosiderin grade per alveolar macrophage or the overall ground truth THS (score for all cells annotated in the slide) were used. The ground truth is a theoretical concept of ‘correct’ annotations.^2^ However, as the ground truth is created by an annotator leading to potential bias, which was tried to mitigate by an computer-assisted labeling approach.

The used ground truth dataset has been published by Marzahl et al. ^22^ and detailed labeling methods can be found in this paper. In summary, development of the final ground truth dataset was done in five consecutive steps: 1) expert-derived annotations of 16 WSIs; 2) development of a deep learning-based algorithm (based on the dataset from step 1); 3) creation of algorithm-derived annotations in the remaining 36 WSIs; 4) diligent review of the expert-derived and algorithm-derived annotations in all 52 WSIs; and 4) review of the assigned label classes assisted by a histogram-like clustering of all annotations. Grading of alveolar macrophages was done according to the definitions by Doucet and Viel (2002). Algorithmic pre-annotations (step 3) of the 36 WSIs was performed to increase efficiency of dataset development (algorithm-expert collaboration). A previous study showed the suitability of this approach for dataset development.^21^ Histogram-like clustering of cell patches based on a continuous, algorithmic regression score (density maps) allows review of the assigned grade and thereby improved consistency of assigning the discrete hemosiderin grades.

### Laboratory evaluation of BALF

To obtain a more objective measure than the mean THS of the annotators or the ground truth THS, we measured other components of blood or its degradation products in BALF, i.e. the RBC count, hemoglobin concentration and iron concentration. Chemical quantification of hemosiderin is not possible. A 5 mL aliquot of BALF was analysed for RBC and hemoglobin content using a CELL-DYN 3700 haematology analyser (Abbott Laboratories, Abbott Park, IL, USA). RBC counts were obtained by means of impedance technology and haemoglobin determination was performed using the modified haemoglobin-hydoxylamine method. The remainder of the fluid aliquot was centrifuged and the supernatant used for iron determination. Iron was measured using an AU680 (Beckman Coulter, Inc, Brea, CA, USA) using a chromogenic method with reduction of iron, and subsequent formation of a complex of ferrous iron with 2,4,6-Tri-(2-pyridil)-5-5triazine which is measured photometrically.

### Statistical analysis

All calculations were performed using R version 4.1.2 (R Foundation, Vienna, Austria). Diagnostic accuracy of EIPH based on the published diagnostic THS cutoff of ≥ 75 ^13^ was calculated in comparison to the ground truth and the mean annotators’ score. To calculate measures of variance including, overall measurement error, error between annotators and residual error, and the intraclass correlation coefficient (ICC) for the THS, a mixed model for a fully crossed, single measure, agreement design (ICC(C,1)) was fitted using the R package lme4 (version 1.1-27.1) (citation). The percentage reduction of overall THS measurement error and reduction in between the annotators due to grade standardization was calculated by fitting two models using the score before and after standardization as outcome.

Cell-level color grading standardization was done by matching the selected cells of every annotator with the cells in the ground truth dataset or the algorithmic dataset, both of which aimed to contain annotations/predictions for every macrophage in the WSIs. Macrophage annotations were considered matching if the Euclidean distance between the center coordinates of both annotations was =< 50 pixels apart. Annotations without a match in the ground truth dataset were excluded from the standardized grading.

To evaluate the effect of the overall measurement error on the diagnosis of EIPH we calculated an 80% uncertainty interval around the Doucet and Viel classification threshold of ≥ 75. This uncertainty interval, analogous to the definition of reference intervals,^14^ is defined as the 10% and 90% quantiles of the annotators’ measurement errors. For individual annotator scores within this interval, the probability that the diagnosis matches the diagnosis of the mean annotators’ score is less than 80% and thus should be considered unreliable and not reproducible.

The correlation between the individual annotators’ scores, the mean annotators’ score, the ground truth score, the algorithmic score, and the BALF RBC count, hemoglobin and iron concentration was calculated using a Spearman correlation.

## Results

All 10 participants annotated at least 300 macrophages in each of the 52 WSIs, thereby creating 158,143 annotations. The ground truth dataset included all alveolar macrophages in the 52 WSIs, and consisted of 215,426 annotations (median: 4,137 per slide; range: 596 to 8,954 per slide). The deep learning-based algorithm analyzed the entire image of the 52 WSIs and detected 218,003 macrophages (median: 3,943 per slide; range: 683 to 8,670 per slide). Time measurements for annotations were available for 7 annotators and 358 WSIs. The median time per case was 14:01 minutes for all annotators combined, and the median time per case ranged between 08:11 to 19:00 minutes for individual annotators. Automated analysis of the entire slides took 1:37 minutes on average (min: 1:31 minutes, max: 1:54 minutes) for each of the 52 WSIs using a modern graphics processing unit (NVIDIA P5000).

### Annotators’ THS: consistency and source of error

The THS had notable variability between the 10 annotators (Fig. 1). The interquartile range of the difference to the mean annotators’ THS was 30 score points (−16 to +14) for all cases combined. The ICC for the THS of the ten annotators was 0.685, i.e. the scoring variance can be explained to 68.5% by a systematic error (difference between annotators in executing the THS) and to 31.5% by a random error (inconsistency within each annotator). Generally, the mean THS of the annotators was somewhat higher than the ground truth THS (on average 25.3 score points). Comparison of the two staining methods revealed that the THS determined from slides stained with modified Turnbull’s blue were somewhat higher than the THS from the corresponding slide stained with Prussian blue (on average 8.5 score points; standard error: 11.1). A somewhat stronger tendency of the THS difference due to the staining methods was observed in the ground truth dataset (average difference of 19.2 score points; standard error of 11.7).

**Fig. 1.**
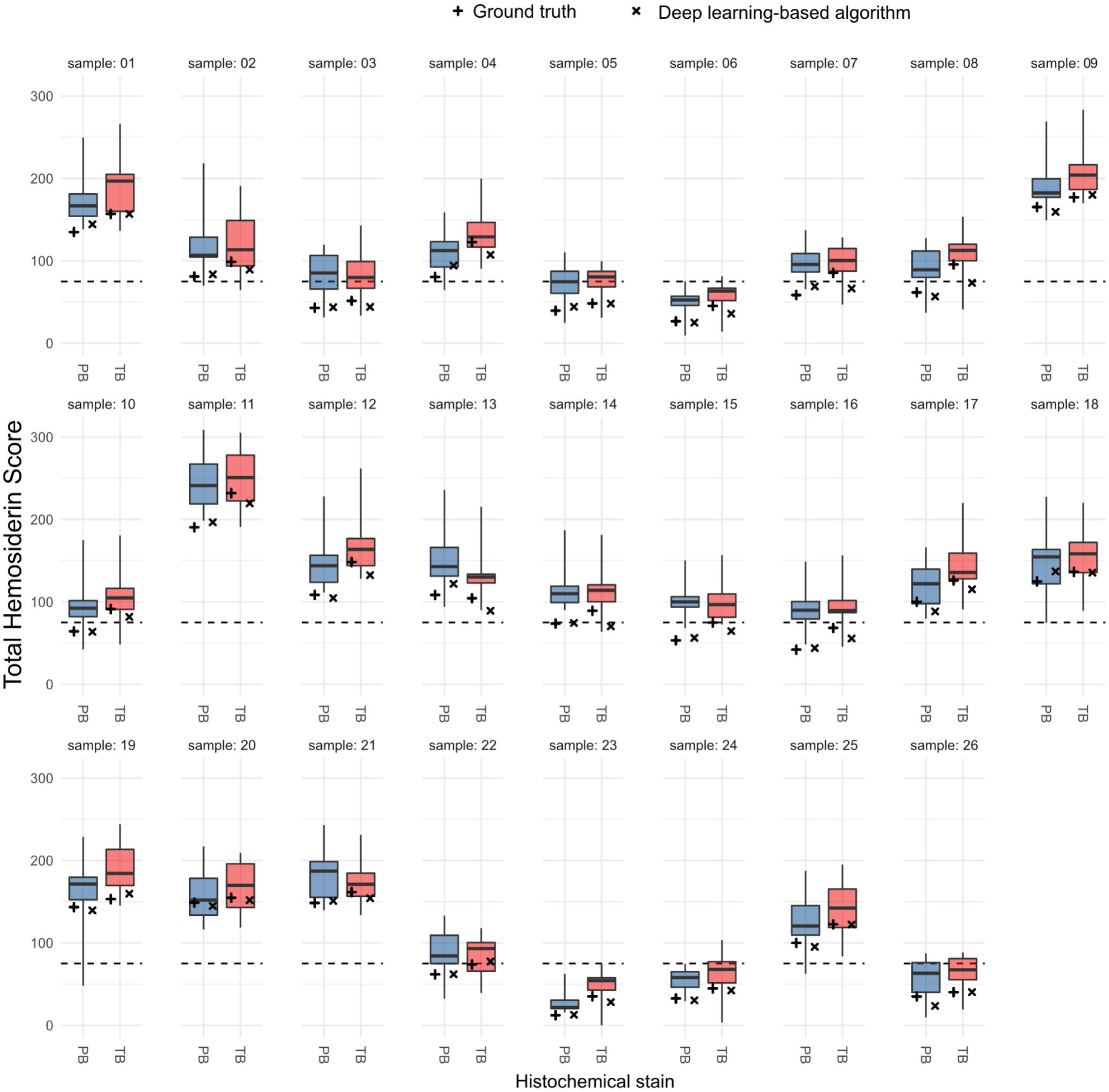
Comparison of the annotators’ total hemosiderin scores (box plots; N = 10) with the ground truth score (+) and the algorithmic score (x) separately for the two staining methods (blue boxplot: Prussian blue, PB; red boxplot: modified Turnbull’s blue, TB). Broken lines represent the cut-off value for diagnosis of exercise-induced pulmonary hemorrhage (total hemosiderin score = 75) published by Doucet et al.^13^

In order to evaluate the variability in hemosiderin grading, we compared the hemosiderin grade of each cell annotated by the annotators with the hemosiderin grade of the ground truth dataset. For the 158,143 annotations by the ten annotators, we could find a cell-matched ground truth label in 121,217 (76.7%) cases. Only 61.7% (76,051 / 121,025) of the matched macrophage annotations had the same hemosiderin grade. Most of the divergent labels (93.6%, 42,411 / 44,974) differed only by one grade level. The annotators assigned a higher grade in 37,919 / 44,974 divergent labels (84.3%). Subsequently, we exchanged the hemosiderin grade assigned by the annotators with the hemosiderin label from the ground truth dataset for all the matched cells, thereby creating a grade-standardized THS. Fig. 2 shows that the measurement error of the THS (difference between the annotators) was markedly reduced when using the standardized hemosiderin grade. Variance analysis determined that the overall measurement error was reduced by 87.8%. The systematic error of annotators was reduced by 97.7%, proving high variability between annotators in applying the published hemosiderin grade stratifications, i.e. judging color saturation. The random error was reduced by 66.4% when the grade-standardized THS was used, which can be explained by the higher consistency in the ground truth dataset that was archived by the multi-step labeling approach (see materials and methods section).

**Fig. 2.**
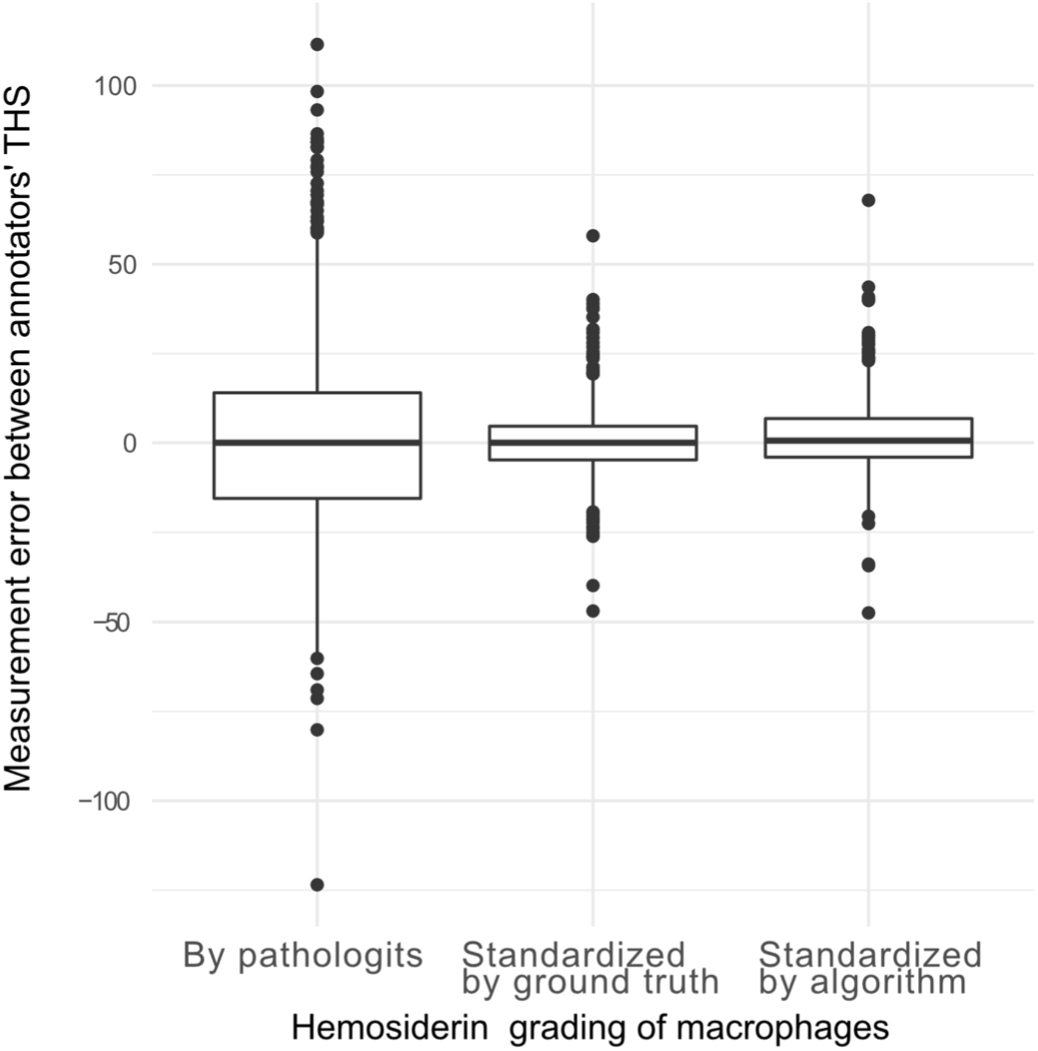
Comparison of the total hemosiderin score (THS) differences of individual annotators to the mean THSs of all annotators. For the left box plot the hemosiderin grade of each included macrophage was derived from the annotators annotations. The measurement error is derived to a large proportion from inter-observer variability in hemosiderin grading (systematic error). For the middle and right box plot the cells selected by the annotators were graded according to the hemosiderin label of the ground truth annotations (middle boxplot) or the algorithmic predictions (right boxplot), thereby eliminating the the inter-observer variability of hemosiderin grading.

### Algorithmic THS: comparison to pathologists’ and ground truth annotations

The algorithmic THS was generally lower than the mean annotators’ THS (on average 28.6 score points), however was quite similar to the ground truth THS (average difference of 3.2 score points; Fig. 1). The mean difference of the algorithmic THS between the two staining methods was 10.6 score points (standard error: 12.7). Algorithmic predictions could be matched (Euclidean distance of ≤ 50 pixels) with 86.6% (186,650 / 218,003) of the ground truth annotations and 76.5% (121,025 / 158,143) of the participants’ annotations. Agreement between the assigned hemosiderin grade labels of these cell-matched annotations was much higher between the algorithm and the ground truth (accuracy: 91.3%; 170,322 / 186,650) than between the algorithm and annotators (62.8%; 76,051 / 121,025). Divergence between the algorithmic and ground truth hemosiderin grades, as well as the algorithmic and annotators’ hemosiderin grades differed mostly by one grade level in 99.9% and 94.3% of instances, respectively. However, annotators had a clear tendency to assign higher grades than the algorithm. Of the cells with divergent hemosiderin grade labels, 84.3% (37,919 / 44,974) of the annotators’ labels were higher than the algorithmic label. In contrast the divergent algorithmic labels had a higher grade level in 50.8% (8,289 / 16,328) and lower grade level in 49.2% (8039 / 8289) of instances, as compared to the ground truth label. This explains why the algorithmic THSs are generally similar to the ground truth THSs but notably lower than the annotators’ THSs.

### Diagnostic accuracy of the annotators’ and algorithmic THS

In 28 of the 52 WSIs (54%), the THS value range of the ten annotators overlaps with the diagnostic cut-off value (THS = 75); thus there would have been inconsistencies in diagnosing EIPH between the ten annotators (Fig. 1). Consensus on the EIPH diagnosis (THS above or below cut-off value) by 8/10 annotators was present in 82.7% of the cases (43 / 52) and consensus by 9/10 annotators was present in 69.2% of the cases (36 / 52). When using the grade-standardized THS of the annotators, the consensus for 9/10 annotators increased to 90.4%.

Compared to the ground truth diagnosis of EIPH (ground truth THS < 75 or ≥ 75), annotators accurately classified the cases in 75.7% with a range of 63.5 - 92.3% for individual annotators (Table 1, Fig. 3). The algorithmic THS had an accuracy of classifying the presence or absence of EIPH of 92.3%. When comparing the algorithmic and individual annotator’s THS to the mean annotators’ THS, diagnostic accuracy was higher for annotators (89.0%) than for the algorithmic approach (71.2%). When analyzing the mean annotators’ THS, the THS range for which there was less than 80% probability of being consistent with a diagnosis of EIPH was 39.2 to 109.8. For 58% instances of the annotator’s THSs that were within this range, the WSIs had not obtained consensus on a diagnosis of EIPH by 9/10 annotators, whereas, in 89% of instances with a THS outside (THS <39.2 or THS > 109.8) of this range the WSIs had achieved consensus in 9/10 annotators.

**Table 1.** Diagnostic accuracy of EIPH (total hemosiderin score above or below diagnostic cut-off of 75) of the total hemosiderin score (THS) of the ten annotators and the deep learning-based algorithm.

|  | Accuracy for diagnosis of EIPH (above or below cut-off) |  |  |  |
| --- | --- | --- | --- | --- |
|  | Compared to the ground truth THS |  |  | Compared to the mean annotators' THS |
|  | All cases (N = 52) | Cases stained with Prussian blue (N = 26) | Cases stained with modified Turnbull blue (N = 26) | All cases (N = 52) |
| All annotators combined | 75.7% | 69.6% | 81.9% | 89.0% |
| Individual annotators | 63.5 – 92.3%<br>(median: 75.0%) | 57.7 - 84.6%<br>(median: 69.2%) | 69.2 - 100%<br>(median: 80.8%) | 60.8 – 98.1%<br>(median: 92.3%) |
| Algorithm | 92.3% | 100% | 84.6% | 71.2% |

**Fig. 3.**
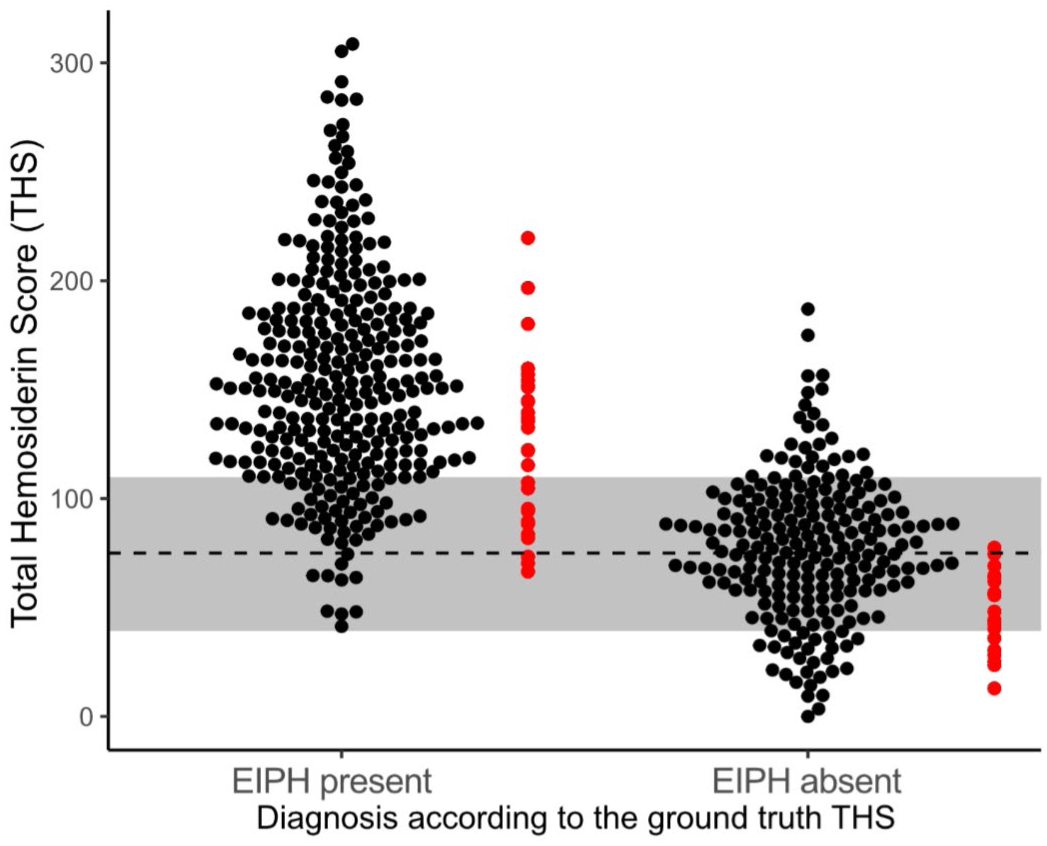
Scatter plots for the total hemosiderin scores (THSs) determined by the ten annotators (black dots) and deep learning-based algorithm (red dots). The 52 cases are separated based on the ground truth THS being above or below the diagnostic cut-off of 75 indicated by the broken line. The grey bar around the broken line is the diagnostic 80% uncertainty interval determined for the human annotators in this study.

### Correlation with the reference methods

The mean annotators’ THSs and algorithmic THSs had very high correlation (R = 0.98) with the ground truth THSs (Table 2). The same correlation (R = 0.98) was identified when the algorithmic THSs were compared with the mean annotators’ THS, whereas individual annotators had a correlation of 0.94 - 0.97 to the mean annotators’ THS.

**Table 2.** Spearman correlation of the annotator’s, ground truth, and algorithmic total hemosiderin score (THS) with the ground truth THS as well as the red blood cell count, hemoglobin concentration, and iron concentration from bronchoalveolar lavage fluid.

|  | Ground truth THS | Red blood cell count | Hemoglobin concentration | Iron concentration |
| --- | --- | --- | --- | --- |
| Mean annotators' THS | 0.98 | 0.08 | 0.11 | 0.75 |
| THS of individual annotators | 0.91 - 0.97<br>(median: 0.94) | -0.11 - 0.14<br>(median: 0.05) | -0.07 - 0.19<br>(median: 0.11) | 0.61 - 0.79<br>(median: 0.72) |
| Ground truth THS | 1.0 | 0.06 | 0.15 | 0.75 |
| Algorithmic THS | 0.98 | 0.12 | 0.20 | 0.79 |

To facilitate an annotator-independent evaluation of the three total hemosiderin scoring methods (the annotators’ THS, the ground truth THS, and the algorithmic THS), these were correlated with the red blood cell count and chemical measurements of hemoglobin and iron concentration. The RBC count and hemoglobin concentration did not correlated with either THS method. The algorithmic THS had a slightly higher correlation with the iron and hemoglobin measurement than the individual or mean annotators’ THS or ground truth THS (Table 2). The iron concentration ranged between <0.4 and 4.7 µmol/L and was below the measurable threshold (<0.4 µmol/L) in nine cases.

## Discussion

The THS by Doucet and Viel (2002) is considered to be one of the most sensitive and accurate tools for the diagnosis of EIPH. However, this method is regarded as too time-consuming for a routine diagnostic test ^9^ and it has not been used in prevalence studies to screen large horse populations. In this study, we evaluated automated image analysis as an approach to improve speed, accuracy, and reproducibility of the THS. Our algorithm was able to score thousands of cells in less than two minutes and had equivalent diagnostic accuracy compared to experts.

In this study, we also evaluated inter-observer variability of the THS and showed that there is high systematic error between experts. Variability between the annotators’ THSs might have resulted from the following sources: 1) bias in selection of the 300 macrophages (representativeness of the included cells), and 2) variability and inconsistency in grading the intracytoplasmatic hemosiderin content of each cell. Regarding the first source of variability, we noticed that the annotators had different selection patterns, which, however, did not seem to have an obvious influence on the participant’s variability. While some annotators screened consecutive fields of view and annotated all macrophages within those fields, others screened the slide in a longitudinal or meandering pattern or selected evenly distributed image locations and annotated some macrophages within these fields (unpublished data).

However, most of the systematic error arose from the differences in applying the hemosiderin grading stratification to alveolar macrophages. Inconsistency in hemosiderin grading was most relevant in cases that were close to the diagnostic cut-off, and led to a lack of consensus by the majority of annotators for the diagnosis of EIPH. For scoring by human experts, we therefore propose to use an 80% uncertainty interval of +/-35 score points around the published cut-off of 75, for which the diagnosis of EIPH is not reproducible by a human expert. Annotators with THS values within this uncertainty interval, i.e. THS values between 40 and 110, had a likelihood of a discrepant diagnosis in more than 20% of cases, when compared to the other annotators, for the presence or absence of EIPH. Our results highlight that increased standardization, or specific training in the application of the scoring system is needed for future studies, or for its use in the diagnostic setting. Development of a standardized “color chart” with images of alveolar macrophages with continuously increasing hemosiderin content and clearly defined thresholds might improve grading consistency between human experts. In our study, we determined that the hemosiderin grading of experts can be standardized by using algorithmic grade predictions, as this led to a marked reduction in the systematic error between annotators.

The present study identified that deep learning-based algorithms are able to achieve high performance for scoring hemosiderophages, that was in many aspects equivalent to the performance of trained experts. However, a major limitation of the present study is the lack of a true gold standard ^9,11,34^ to compare the performance of expert annotators with the performance of the deep learning-based algorithm without bias. For the development of deep learning-based algorithms for histopathological and cytological tasks, it is often the gold standard to compare the algorithmic predictions with expert-derived ground truth annotations.^2,5,20,24^ Nevertheless, it needs to be acknowledged that human errors in the ground truth labeling may have a bias on performance evaluation. This is why we sought to mitigate human errors in the ground truth dataset by using a multi-step, computer-assisted labeling approach,^22^ which our research group validated for this specific task in previous studies.^21^ As this ground truth dataset were also used to train the algorithmic models (using a three-fold cross validation), we found very high consistency between the ground truth dataset and algorithmic predictions, indicating that the models replicated the training data very well. When the ground truth THSs were used as the reference, the algorithm had a higher diagnostic accuracy than the annotators. In contrast, the ten annotators had the clear tendency to assign higher hemosiderin grades to alveolar macrophages, i.e. systematically applied lower thresholds for the individual hemosiderin grades than the ground truth annotator and algorithm. This explains the marked difference between the THSs of the study participants and the ground truth and algorithmic individual THSs as well as the higher diagnostic accuracy of the individual annotator’s THSs compared to the mean annotators’ THS.

Due to the above mentioned bias of the annotators’ THS and ground truth dataset as a reference method, we evalauted three observer-independent reference methods (RBC, hemoglobin and iron concentration in BALF). We found that the iron concentration had a high correlation with the THSs, whereas the RBC and hemoglobin concentration did not correlate with the THSs. RBC and hemoglobin concentrations are features of acute pulmonary bleeding and are degraded shortly after the hemorrhagic event and therefore seem to be inappropriate reference methods for the THS, which measures chronic hemorrhage. In the present study, we used iron concentration for the first time as a measure of pulmonary bleeding. The limitation of the chemical iron measurement was the low iron content in BALF which in some cases was below the limit of detection. Future studies are needed to determine the value of iron concentration as a reference method for the THS and as a potential diagnostic test for EIPH.

Automated image analysis using deep learning is a highly relevant field of research in veterinary clinical, anatomic and toxicologic pathology that is generally considered promising for increasing accuracy, reproducibility and time efficiency for quantitative tasks.^1,5,6,23,27,35^ A precondition for computerized analysis is the availability of digital images, which is aided by the current trend of digitizing the diagnostic workflow of pathology laboratories. The use of digital microscopy for cytological specimens is, however, hampered by limited image resolution and lack of fine focus of default WSIs.^6,7^ Nevertheless, the study participants consider WSIs appropriate to perform the THS in this study’s cases, because of the uniform depth of the samples, and as relatively little cellular detail is necessary to evaluate the intracytoplasmic hemosiderin content of macrophages. Another limitation of WSIs specific to the task of scoring hemosiderophages is that different WSI scanners often exhibit a marked difference in the color representation, i.e. might have a higher or lower intensity of the blue color. This is likely to influence annotators and algorithms in evaluating the amount of blue pigment and needs to be evaluated in future studies. Currently there are few studies that have evaluated the benefits of automated image analysis compared to the visual assessment by experts in veterinary medicine.^4,5,8,20^ These studies are needed to critically evaluate potential sources of algorithmic errors before an algorithm can be used for routine diagnostic purposes. Future studies need to evaluate how THS algorithms are best implemented in a diagnostic workflow, while ensuring high diagnostic reliability. Generally, algorithms can be used to automatically predict the diagnosis or they can be used as an assistive tool, which supports annotators in critical steps of the diagnostic task (computer-assisted diagnosis). For the diagnosis of EIPH, THSs could be derived fully automatically by the algorithm, with only rough verification of the predictions by an expert. Alternatively, algorithms could be used to standardize hemosiderin grading of individual macrophages that are selected by annotators (computer-assisted THS). The benefits of both applications on diagnostic accuracy and reproducibility have been demonstrated in this study.

## Conclusion

Cytologic quantification of hemosiderophages in BALF using the THS is considered the most sensitive method for the diagnosis of EIPH. However, we have shown that the THS by human experts is time-consuming and has a high systematic error between observers. We propose to use an uncertainty interval of 75 +/-35 score points for the diagnosis of EIPH by experts. Furthermore, to overcome the limitations of human experts, we validated a deep learning-based image analysis algorithm that had high accuracy/correlation compared to the mean THSs of ten annotators, a ground truth dataset and iron concentration of BALF. We have shown that deep learning-based algorithms are a valuable tool for time-efficient, accurate, and reproducible scoring of hemosiderophages, which could be applied to research studies, such as large prevalence studies, and routine diagnostic service.

## Acknowledgement

C.A.B. gratefully acknowledges financial support received from the Dres. Jutta and Georg Bruns-Stifung für innovative Veterinärmedizin.

